# Understanding the development of oral epithelial organs through single cell transcriptomic analysis

**DOI:** 10.1101/2022.01.18.476858

**Authors:** Qianlin Ye, Arshia Bhojwani, Jimmy K. Hu

## Abstract

During vertebrate craniofacial development, the oral epithelium begins as a simple and morphologically homogeneous tissue. It then gives rise to locally complex structures, including the developing teeth, salivary glands, and taste buds. While there is significant knowledge about the molecular mechanisms regulating the morphogenesis of these organs at later stages, how the epithelium is initially patterned and specified to generate diverse cell types and organs remains largely unknown. To elucidate the genetic programs that direct the formation of distinct oral epithelial populations, we mapped the transcriptional landscape of embryonic day (E) 12 mouse mandibular epithelia at single cell resolution. Our analysis identified key transcription factors and gene regulatory networks that define different epithelial cell types as well as regions patterned along the oral-aboral axis. By examining the spatiotemporal expression of region-specific markers in embryonic mandibles, our results pointed to a model where the dental field is progressively confined to its position by the formation of the aboral epithelium anteriorly and the non-dental oral epithelium posteriorly. Using our data, we also identified *Ntrk2* as a promoter of cell proliferation in the forming incisor, contributing to its invagination. Together, our results provide a detailed transcriptional atlas of the developing mandibular epithelium and unveil new genetic markers and regulators that are present during the specification of various oral epithelial structures.

## Introduction

The vertebrate mouth is a highly derived and complex structure that is crucial for the survival of animals [1–3]. It consists of specialized organs – teeth, taste buds, and glands – that are part of the feeding apparatus to attain, taste, moisten, and ingest food. During development, these organs emerge as epithelial thickenings or placodes at precise locations on the oral surface of the first pharyngeal arch and then invaginate into the underlying mesenchyme as they undergo morphogenesis [4–6]. The sculpting of the oral epithelium from an initial monolayer into a stratified tissue with distinct structures and organ types requires spatiotemporal patterning of the epithelium. This imparts lineage-specific transcriptional changes for the correct specification of different epithelial cell fates.

The mouse mandible, which constitutes the lower part of the mouth, has served as one of the foundational model systems to study the development of oral epithelial organs [7]. Like in all gnathostomes, or jawed vertebrates, the mouse jaw is formed from the mandibular process of the first pharyngeal arch which comprises a mesenchymal core ensheathed by a contiguous epithelial layer of both ectodermal and endodermal origins [8]. In mice, the ectoderm covers the outer mandible and most of the oral surface, while the endoderm overlays the inner mandible that extends posteriorly from the proximal part of the tongue to the embryonic foregut [9]. Patterning of the mandibular epithelium is already evident at E9.5, shortly after the mandibular process forms. *Bmp4* and *Fgf8* are expressed in the medial (distal) and lateral (proximal) portion of the oral epithelium respectively at this stage, and their encoded ligands signal to both the epithelium and the underlying mesenchyme to pattern the incisor and molar fields along the proximal-distal axis [10–12]. Concurrently, *Shh* is expressed in the pharyngeal endoderm and signaling by SHH and BMP4 patterns the mandible along the oral-aboral axis [13]. These signaling events thus organize the mandible into a few broadly patterned domains prior to the formation of morphologically distinct epithelial placodes.

The oral epithelium begins as a single layer of cuboidal cells but soon undergoes stratification between E10.0 and E11.0 to produce a sheet of flattened cells called the periderm that coats the epithelia on the apical side and protects them from inappropriate fusion [14]. Further stratification in the developing ectodermal organs generates suprabasal cells that stack between the columnar basal layer and the periderm. In the case of the developing teeth, this thickened epithelium, known as the dental lamina, is discernible as a horseshoe-shaped stripe across the oral surface at E11.0 and represents the earliest morphological sign of tooth formation [15,16]. However, the exact mechanism that determines the position of the dental lamina and initiates tooth development remains unclear. Cells in the dental lamina are transcriptionally distinct from the rest of the epithelium, expressing several tooth-specific transcription factor genes, including *Pitx2*, *Irx1*, and *Foxi3* [17–19]. In particular, *Pitx2* is critical for tooth development and is one of the earliest dental markers [20]. Its expression precedes the lamina stage and is present in a broader domain at E10.5 that later narrows and localizes to the invaginating dental epithelium [21]. Patterning of the dental field is therefore a progressive process involving dynamic transcriptional changes.

As the dental lamina transitions through the placode and bud stages between E11.0 and E12.5, the early tooth signaling center, called the initiation knot, can be identified by its expression of *Shh*, *Bmp2*, *Fgf20*, and *Wnt10a/b* along the anterior tooth epithelium [22–24]. Signaling from the initiation knot plays a key role in promoting proliferation of dental epithelial cells, controlling the size of the tooth bud, and driving its invagination [25–28]. Thus, by E12.0 individual incisor and molar buds are easily recognizable. They are separated by a toothless space called the diastema, as mouse dentition is reduced and lacks canines and pre-molars. Besides teeth, other oral ectodermal organs also begin to form around this stage. For example, the development of the submandibular salivary gland initiates at E11.5, first as an epithelial thickening adjacent to the tongue, which then protrudes into the mesenchyme as a teardrop shaped bud by E12.0 [29,30]. Concomitantly, taste bud primordia develop as taste placodes on the mouse tongue starting at E12.0 [31,32]. Therefore, E12.0 represents a critical developmental window for experimental investigations, as it is marked by the emergence and expansion of different progenitor lineages that will form all the major oral ectodermal organs. Crucially, the exact mechanisms by which diverse mandibular epithelial populations are patterned and specified remain an important open question. As many of the studies to date have focused primarily on individual organs or other timepoints [33–38,13], there is also incomplete knowledge in the overall cell heterogeneity within the E12.0 mandibular epithelium, as well as how specific gene regulatory networks and signaling processes are coordinated across both space and time to control the development and functions of different epithelial populations.

To address these questions, we first acquired an in-depth understanding of the cell diversity in the mandibular epithelium at E12.0 using single cell RNA-sequencing (scRNA-seq), which enables transcriptomic profiling of thousands of individual cells simultaneously [39,40]. By complementing sequencing results with detailed spatial mapping using RNA *in situ* hybridization, we have identified discrete populations not only in the developing ectodermal organs but also in areas where the epithelium appears morphologically simple and uniform. Through computational analysis, we uncovered key transcription factors and associated gene regulatory networks that define these epithelial regions and cell lineages. In addition, we show that the oral-aboral patterning of the mandibular epithelium evolves over time as it becomes increasingly regionalized at the transcriptional level from E9.5 to E12.0. Our findings indicate that the mandibular ectoderm is initially more dental-like with the expression of tooth-specific transcription factors, but newly patterned anterior and posterior regions gradually expand and confine the dental progenitor field to its eventual position. Lastly, we show that *Ntrk2*, a novel dental marker we identified in our analysis, promotes cell proliferation and tooth invagination. Our results thus provide a useful resource for future investigation of key regulators during mandibular epithelial morphogenesis.

## Methods

### Mouse lines and colony maintenance

*K14^Cre^* [41], *R26^mT/mG^* [42], *Shh^CreER^* [43] and *Tagln^Cre^* mice[44] were group housed and genotyped as previously published (sequences provided in Table S1). Except *K14^Cre^*, all mice were acquired from the Jackson Laboratory (JAX) and maintained on a C57BL/6J background. *K14^Cre^* was on a mixed background at the time of acquisition but subsequently crossed to *R26^mT/mG^* (C57BL/6J) for more than 6 generations. The resulting *K14^Cre^*;*R26^mT/mG^* mice were used to produce embryos for the scRNA-seq, RNA *in situ* hybridization mapping, and explant culture experiments in this study. For lineage tracing, *Shh^CreER^* and *Tagln^Cre^* were crossed to *R26^mT/mG^* to produce *Shh^CreER^*;*R26^mT/mG^* and *Tagln^Cre^*;*R26^mT/mG^* respectively. Timed pregnancy was set up either in the morning or in the afternoon to obtain embryos at different stages as indicated in the text. Noon of the day of vaginal plug discovery was designated as E0.0 or E0.5 depending on the time of breeding setup. Both male and female embryos were selected randomly and used in all experiments. To activate CreER, tamoxifen dissolved in corn oil at a dose of 1.25 mg/30 g body weight was delivered to pregnant *Shh^CreER^*;*R26^mT/mG^* females at E8.5 and E9.5 through oral gavage. All mice were maintained in the University of California Los Angeles (UCLA) pathogen-free animal facility. All experiments involving mice were approved by the Institutional Animal Care and Use Committee of UCLA (Protocol Number ARC-2019-013.

### Single cell isolation from mouse embryonic mandibles

The protocol for single cell dissociation was modified from previous studies [45,46]. To isolate mandibular epithelial cells for scRNA-seq, we harvested six E12.0 *K14^Cre^;R26^mT/mG^* mouse embryos from the same litter and dissected out their mandibles in cold HBSS (Gibco). Mandibles were then pooled and incubated with 10 mg/ml Dispase II (Sigma-Aldrich) in HBSS supplemented with 10 µg/ml DNase (New England Biolabs) at 37°C and swirled at 100 rpm for 32 minutes to enzymatically separate the epithelium from the mesenchyme as we previously did [47]. After peeling off the epithelia using forceps, they were dissociated in TrypLE (Gibco) at 37°C for 30 minutes, with gentle pipetting every 10 minutes. Cells were then centrifuged at 400 rcf for 5 minutes and resuspended in cold flow cytometry buffer (calcium free HBSS with 5% fetal bovine serum (FBS, Gibco), 2 mM EDTA, and 10 mM HEPES (Gibco)). Undissociated cell clumps were sieved out using a 20 µm pluriStrainer and the resulting cell suspension was sorted using fluorescence-activated cell sorting (FACS) to isolate GFP+ single epithelial cells. Cell numbers and viability were analyzed using the Invitrogen Countess II FL, which showed greater than 90% viability.

### Single-cell RNA-seq: barcoding, library construction, and data analysis

The live single cell numbers in suspension were adjusted to a final concentration of 1000 cells/µl in PBS with 0.04% BSA and approximately 14000 cells were loaded to a 10X Chromium Single Cell instrument for single cell partitioning at the UCLA Technology Center for Genomics and Bioinformatics (TCGB). Sample barcoding, cDNA amplification and library construction were performed using the Chromium Single Cell 3ʹ Library Kit v3 according to the manufacturer’s instructions. The cDNA library was confirmed for its quality using an Agilent TapeStation. A total of 11,131 cells were successfully sequenced on a Illumina NextSeq 500 system, which produced about 218 million reads.

The sequencing reads were aligned against GRCm38 using CellRanger 2.0.0. Further downstream analyses were conducted in the R package Seurat [48]. Following the standard Seurat workflow, we first filtered out low quality cells with less than 200 or over 5500 unique feature counts or with more than 10% mitochondrial gene counts. The filtered dataset was then normalized using Seurat’s SCTransform function [49]. To reduce the effects of cell cycle and sex heterogeneity in our scRNA-seq data, we first used Seurat’s CellCycleScoring or AddModuleScore functions to assign scores to these categories based on a list of cell cycle genes (G2/M and S markers) [50] and sexually dimorphic genes (*Uty*, *Ddx3y*, *Kdm5d*, *Eif2s3y*, *Xist*, *Tsix*, and *Lars2*) [51,52], which were then regressed out from the count matrix as described in the Seurat documentation. We have also regressed out genes encoding lincRNAs, Gm42418 and AY036118, which overlap the rRNA Rn45s locus and can be differentially amplified as an artefact at the amplification step [53].

We next performed dimensionality reduction by Principal Component Analysis (PCA) and Uniform Manifold Approximation and Projection (UMAP), followed by unsupervised cell clustering using the FindNeighbors and FindClusters functions. The number of top principal components (PCs) used for dimensional reduction and the resolution of clustering were guided by Seurat’s ElbowPlot and the Clustree package [54] respectively. This allowed us to first obtain an overview of the general cell populations using a low-resolution parameter (10 PCs and 0.08 resolution) and then examine the constituent sub-populations in greater detail with a high-resolution setting (30 PCs and 0.9 resolution). For the peridermal and tongue epithelium clusters, we further subset these cells and iterated clustering to identify the different cell types within. Differentially expressed marker genes for each cluster were identified by Seurat’s FindAllMarkers function using the Wilcoxon Rank Sum test, with the cutoff criteria set for genes expressed in a minimum of 15% of cells and a fold change of 1.3 (Table S2). Functional enrichment analysis for top ranked cluster(s)-specific marker genes with adjusted P-value p_val_adj ≤ 1 x 10-50 was performed using Metascape (http://metascape.org) [55]. As most marker genes in clusters Di are not highly specific and also enriched in neighboring clusters, we did not include its functional enrichment analysis in this study. Differentially expressed genes with a pct. 2 < 0.5 were used to assess the number of markers co-expressed by cells in the dental, taste bud, and salivary gland clusters.

The E10.5 mandibular epithelial scRNA-seq data were subset from a previously published whole mandible single-cell dataset [13], pre-processed with SCTransform and regression of effects from cell cycle, sex, and lincRNAs, and then integrated into our dataset using Seurat as previously described [56]. Slingshot pseudotime lineage inference was applied through the Dynverse package [57,58]. To identify differentially expressed key transcription factors and their downstream gene regulatory networks, we applied SCENIC (single-cell regulatory network inference and clustering) to analyze our data using the default setting [59]. To better visualize the network, we also examined marker genes from each or combined clusters using the iRegulon plugin from Cytoscape [60], setting the putative regulatory region at 500 bp upstream and an enrichment score threshold of 2.5.

### RNA *in situ* hybridization

Mouse embryos at different stages were dissected out from the uterus in DEPC-treated PBS. For whole mount RNA *in situ* hybridization (WISH), the mandibles were collected and fixed with 4% paraformaldehyde (PFA) in DEPC-treated PBS overnight at 4°C. WISH was carried out as previously described [61]. For each marker gene, we designed two anti-sense digoxigenin-labeled probes whenever possible, unless restricted by gene size, sequence homology, or cloning challenges (Table S1). Hybridized tissues were detected by BM Purple (Roche) and imaged using a Leica DFC7000 T camera fitted on a Leica M205 stereomicroscope. To further analyze gene expression in the epithelium at a finer resolution, the stained whole mount samples were processed through serial sucrose washes and embedded in the Tissue-Tek O.C.T. compound (Sakura Finetek) for frozen sections. 10 μm thick sections were obtained using a Leica CM3050S Cryostat and imaged using a Leica DM 1000 microscope. For samples with weak signals following sectioning, we instead performed section RNA *in situ* hybridization on paraffin sections (7 μm) using established protocols [62].

For RNAscope analysis, embryonic heads were dissected and fixed in 10% neutral buffered formalin for 24 hours at room temperature and dehydrated through serial ethanol washes, embedded in paraffin, and sectioned at 6 μm. RNAscope was carried out using the RNAscope Multiplex Fluorescent v2 Assay (ACD) by following the manufacturer’s instructions. Optimized tissue pretreatment steps include boiling sections in the Target Retrieval Reagents (ACD) at 100°C for 10 minutes and incubating samples in the Protease Plus solution (ACD) at 40°C for 10 minutes. Opal 520, 570, and 690 from Akoya Biosciences were used for color development. RNAscope 3-plex Negative Control Probe (ACD) consistently showed no background staining. RNAscope *Mus musculus* probes *Bdnf, Cxcl14*, *Ddit4l*, *Dmrt2, Irx1*, *Ntrk2*, *Pitx2*, *Rxfp1, Shh*, *Tfap2b*, and *Zcchc5* were purchased from ACD. The *Pitx2* probe set recognizes all isoforms, which were shown to have similar expression patterns at the stages examined in this study [20].

### Immunofluorescence staining

For Immunofluorescence staining, samples were fixed in 4% PFA overnight at 4°C and prepared for either frozen or paraffin sections. For paraffin sections, antigen retrieval was performed by sub-boiling slides in a microwave for 15 minutes in citric buffer (pH 6.2) containing 10 mM citric acid, 2 mM EDTA, and 0.05% Tween-20. After blocking tissues with a blocking solution (1X animal-free blocker (Vector Laboratories), 2% heat inactivated goat serum, 0.02% SDS, and 0.1% Triton-X) for 1 hour, slides were incubated with the following primary antibodies overnight at 4°C: ACTA2 (Abcam, ab8211), E-cadherin (Cell Signaling, 3195S), and GFP (Abcam, ab13970). All antibodies were diluted at 1:100 in the same block without serum. Secondary antibodies (Thermo Scientific) used include Alexa Fluor 555 and Alexa Fluor 488 goat anti-rabbit IgG, Alexa Fluor 488 goat anti-mouse IgG, and Alexa Fluor 488 goat anti-chick IgG, all at 1:250 dilution for 1 hour at room temperature. DAPI (Invitrogen) was used as a nuclear stain. 5-ethynyl-2′-deoxyuridine (EdU) incorporation was detected using a Click-iT Plus EdU Alexa Fluor 555 Assay Kit (Invitrogen, C10638) prior to primary antibody incubation. All images were taken using a Zeiss LSM 780 confocal microscope.

### Explant culture

Dissected E11.5 embryonic mandibles were cultured on top of a 0.4 μm Millicell filter (Millipore) supported by a metal mesh (914 μm mesh opening, Spectrum Labs) at the interface of air and media at 37°C and 5% CO_2_ as previously reported [63]. The culture media contains BGJb medium (Gibco), 3% FBS (Gibco), 1% MEM non-essential amino acids (Gibco), 1% GlutaMax (Gibco), 140 μg/ml L-ascorbic acid (Thermo), 1% penicillin-streptomycin (Thermo), and with the NTRK2 inhibitor ANA-12 (150 μM, Sigma-Aldrich) or equal volume of DMSO control vehicle. The mandible explants were cultured for either 48 or 72 hours before processing for paraffin sections. For labelling cycling cells with EdU, 5 μl of EdU (10mg/ml, Thermo Scientific) was directly pipetted on top of the explants and then incubated for 4 hours before processing tissues for frozen sections. The 4 hours pulse time was optimized to ensure sufficient labelling, as shorter pulses marked fewer cells due to slower explant development. Quantification of the tooth germ size was carried out using ImageJ.

### Statistics and reproducibility

All experiments, except scRNA-seq and RNA *in situ* hybridization, were replicated at least three times using independent biological samples. Marker genes identified from scRNA-seq were validated by RNA *in situ* hybridization studies, which were conducted in at least two independent biological experiments per probe. RNAscope was replicated in at least 3 independent biological experiments for each set of double Z oligo probes. All images are representative. Each data point in Figs. 7M-O and 9B,D represents a single biological sample. Data points were collected without investigator blinding. No data were excluded. Graphs were prepared using the Prism software and display mean ± s.d. (standard deviation). *P-*values were calculated as specified in figure legends. Significance was taken as *p* < 0.05 with a confidence interval of 95%. * *p* < 0.05; ** *p* < 0.01; *** *p* < 0.001; **** *p* < 0.0001.

## Results

### scRNA-seq identifies spatially distinct epithelial populations in the developing mandible

In order to define the different epithelial populations in the developing mandible based on their genetic differences, we performed scRNA-seq analysis using mandibular epithelia dissected from E12.0 embryos. At this timepoint, the oral epithelium has been broadly patterned along the oral-aboral and medial-lateral axes (Fig. 1A) [13], and several epithelial organs, such as the tooth and the salivary gland, have just begun to develop and actively undergo stratification and invagination. The E12.0 mandible therefore provides an ideal platform to investigate the intrinsic genetic regulation governing the development and functions of all the early epithelial progenitor populations that constitute each of the mandibular ectodermal structures and their adjoining regions. To label the epithelium for easier downstream processing, we utilized *K14^Cre^;R26^mT/mG^* embryos, where Keratin 14-driven Cre recombinase permanently labels epithelial cells with membrane GFP (mG) from the *R26^mT/mG^* Cre reporter. Mesenchymal cells lack Cre activity and continue to express membrane tdTomato (mT). We first enzymatically separated the epithelium from the mesenchyme before dissociating the epithelium alone into a cell suspension (Fig. 1B). This allows a more uniform dissociation process and capture of most epithelial cells, which are more adhesive than the mesenchyme and far fewer in number comparatively. As cell doublets can confound scRNA-seq data interpretation [64], we next performed FACS to sort GFP+ single epithelial cells (Fig. 1C). After barcoding cells using the 10X Chromium Controller (Fig. 1D), we sequenced a total of 11,131 cells from 6 mandibles and visualized the unsupervised clustering of the scRNA-seq data using UMAP under the Seurat package [48]. The initial feature plot contains mirrored clusters that differ in the expression of cell cycle related genes, such as *Ccnb1* and *Top2a* (Fig. S1), reflecting the proliferative nature of a developing tissue. To reduce the clustering complexity and to focus on the transcriptional differences associated with epithelial sub-structures, we subsequently regressed out transcripts related to cell cycles as well as transcripts segregated with sexes.

**Figure 1.**
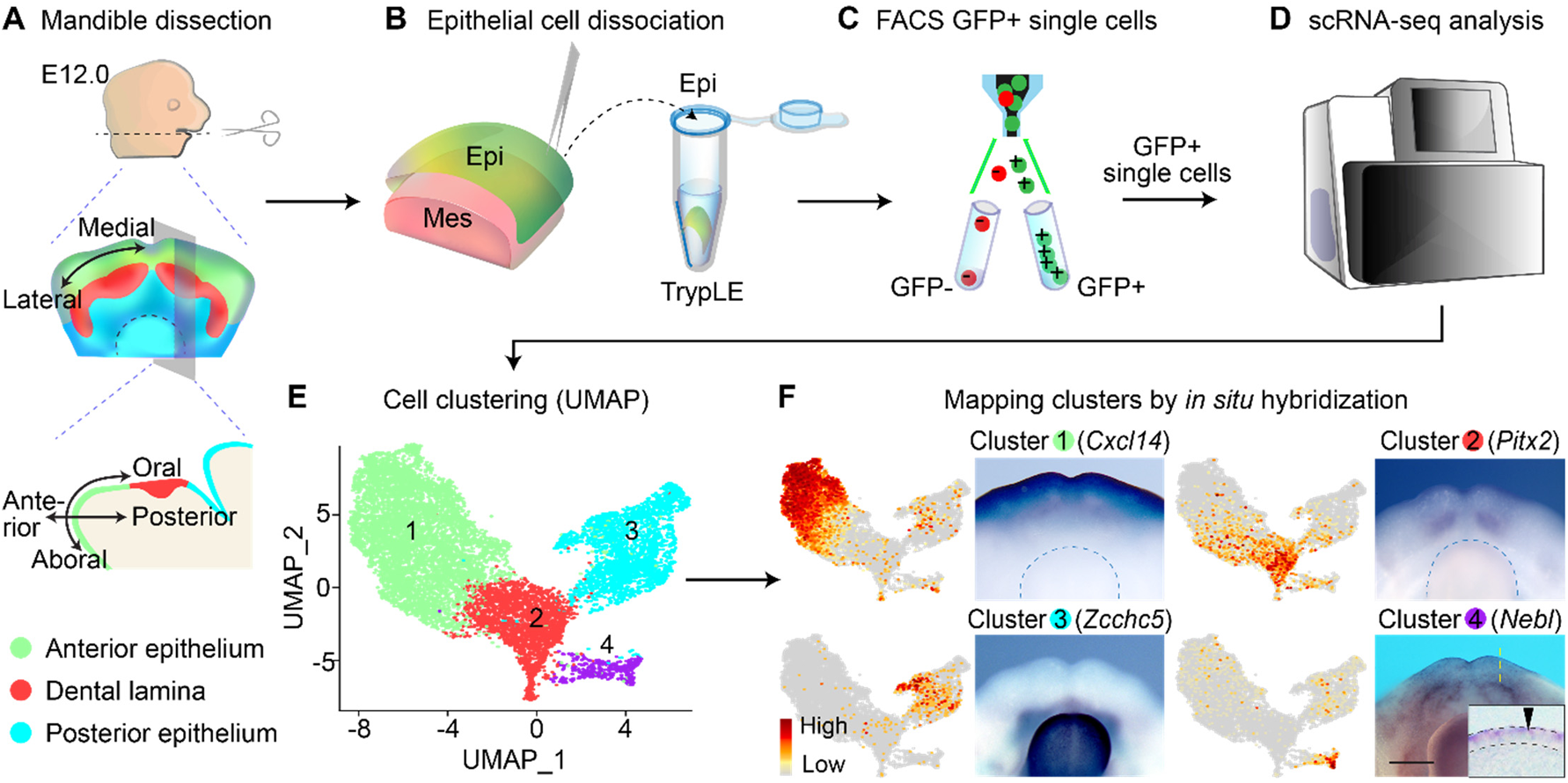
Single cell RNA-sequencing of E12.0 mandibular epithelium. (**A-D**) Outline of the workflow for cell isolation and single-cell transcriptome profiling of E12.0 mandibular epithelia. Schematic drawings of the embryonic mouse mandible in dorsal view and sagittal section showing the anatomical axes and broadly defined epithelial regions. (**E**) UMAP plot of mandibular epithelial cells showing four major clusters. (**F**) Feature plots of representative marker genes enriched in the four clusters shown in (E) and their expression by RNA *in situ* hybridization in E12.0 mandibles (dorsal views). Clusters 1-3 correspond to the anterior (green), dental (red), and posterior (cyan) epithelium respectively. Blue dashed lines outline the tongue. Cluster 4 (purple) contains periderm cells (black arrowhead). Inset is a representative sagittal section through the yellow dashed line. Black dashed lines demarcate the epithelium. Epi, epithelium; Mes, mesenchyme. Scale bar represents 400 μm in whole mount images and 20 μm in the inset.

We first analyzed this refined dataset at a low resolution, and this partitioned the mandibular epithelium into 4 large clusters (Fig. 1E). Cluster 1 differentially expresses several markers, including *Cxcl14* and *Tfap2b*. To locate these cells in the E12.0 mandible, we performed whole mount RNA *in situ* hybridization using cluster 1 markers and mapped them to the aboral epithelium that is anterior to the dental lamina and continues to the ventral mandible (Fig. 1F, Fig. S2A). This result also corroborates the *Tfap2b* expression pattern previously reported [65,66]. Cluster 2 expresses many known dental markers, such as *Pitx2* [67] and *Irx2* [68] (Fig. 1F, Fig. S2B), and thus contains cells from the developing teeth. Cluster 3 could therefore in theory occupy the region posterior to the dental lamina. Analysis of cluster 3 unveiled many previously unidentified markers of the oral epithelium, including *Zcchc5* and *Col14a1* (Fig. 1F, Fig. S2C). Examination of their expression verified that cluster 3 represents the epithelium posterior to the forming teeth. The positions of clusters 1-3 on the UMAP from left to right therefore correspond spatially to the mandibular epithelium from the ventral and anterior aboral domain to the posterior oral surface that includes the tongue (Fig. 1A,E). Cluster 4 is defined by the expression of several known periderm markers, including *Grhl3*, *Irf6*, and *Sfn* [69–71]. The periderm is a layer of flattened cells that coats the developing epithelia and analyzing the expression of other cluster 4-specific markers, such as *Nebl* and *Pkp1*, confirmed the inclusion of periderm in this cluster (Fig. 1F, Fig. S2D).

To obtain a more detailed classification of the different sub-populations in the mandibular epithelium, we re-clustered the cells at a higher resolution. This yielded 15 clusters that are characterized by distinct gene expression signatures (Fig. 2). By identifying marker genes enriched in each cluster and mapping their spatial distribution within the epithelium, we were able to assign the identities of each cluster according to their anatomical positions. These are V1 (ventral 1), V2 (ventral 2), AVM (anteroventral-medial), AVL (anteroventral-lateral), ADM (anterodorsal-medial), and ADL (anterodorsal-lateral), which constitute cluster 1 described above; Di (diastema), In (incisor), Mo (molar), and IK (initiation knot), which subdivide cluster 2; PM (posterior-medial), PL (posterior-lateral), SG (salivary gland), and T (tongue) from cluster 3, and P/S (periderm and suprabasal cells) that makes up cluster 4. We further describe these clusters and their associated markers below.

**Figure 2.**
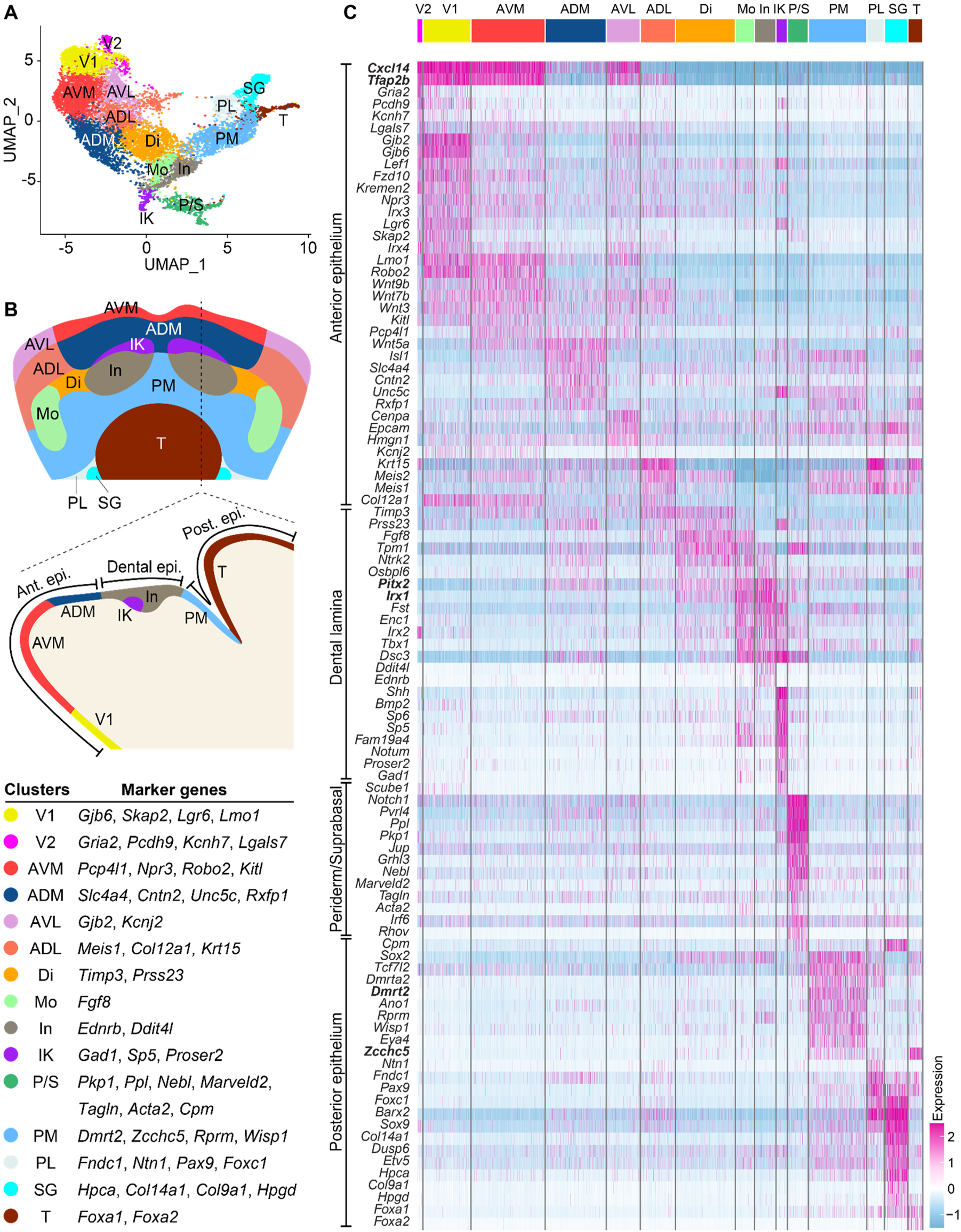
Mandibular epithelial cells are clustered based on their anatomical positions. (**A**) Second level UMAP clustering reveals 15 epithelial cell clusters. (**B**) Schematic drawings of the mandible in dorsal view and sagittal section showing the anatomical positions of the 15 clusters based on RNA *in situ* mapping, which partition the anterior (ant.), dental, and posterior (post.) epithelium (epi.) into subdomains. Representative marker genes used for *in situ* hybridization to assign clusters are shown. (**C**) Heatmap of differentially expressed genes in each cluster showing scaled expression level from low (blue) to high (pink). Genes in bold are markers for the broader epithelial regions indicated on the left. Clusters assigned: V1, ventral 1; V2, ventral 2; AVM, anteroventral-medial; ADM, anterodorsal-medial; AVL, anteroventral-lateral; ADL, anterodorsal-lateral; Di, diastema; Mo, molar; In, incisor; IK, initiation knot; P/S, periderm and suprabasal cells; PM, posterior-medial; PL, posterior-lateral; SG, salivary gland; T, tongue.

### Anterior mandibular epithelium is patterned into subdomains and expresses regulators of WNT and BMP pathways

We first examined clusters corresponding to cluster 1 above, the epithelial region that extends from the ventral aboral side of the mandibular epithelium and ends anterior to the dental lamina (Fig. 1A,E). Cluster V1 includes cells that occupy the ventral mandibular epithelium, based on the expression of its markers, *Gjb6*, *Skap2*, *Lgr6*, and *Lmo1* (Fig. 3A, Fig. S3A-C). V2 shares a similar expression profile as V1 but also differentially expresses a set of genes that include *Gria2*, *Kcnh7, Lgals7,* and *Pcdh9* (Fig. 2C). Expression analysis of these transcripts found that V2 cells were dispersed as puncta within the ventral epithelium, as well as expressed in two lateral epithelial patches on either side of the ventral mandible (Fig. 3B, Fig. S3D-F). We next studied cells from clusters AVM, ADM, AVL, and ADL. Probing the expression of markers for AVM (*Pcp4l1*, *Npr3*, *Kitl*, and *Robo2*) and ADM (*Slc4a4*, *Cntn2*, *Rxfp1*, and *Unc5c*) showed that these two clusters contain cells in the medial portion of the mandible that is anterior to the forming incisor (Fig. 3C,D, Fig. S3G-L). AVM cells are located in between ADM and V1 cells, consistent with their relative positions on the UMAP (Fig. 2A). Adjacent to the AVM and ADM clusters are AVL and ADL. While the pairs of AVM/AVL and ADM/ADL share many transcriptional features (Fig. 2C), assessment of genes differentially expressed in AVL (*Gjb2* and *Kcnj2*) or ADL (*Meis1*, *Col12a1*and *Krt15*) but reduced in AVM/ADM (Fig. 3E,F, Fig. S3M-O) indicates that AVL and ADL cells are positioned in the lateral portion of the anterior mandible (Fig. 3G). Therefore, while the x-axis of the UMAP corresponds to the oral-aboral axis of the mandible, the y-axis matches the medial-lateral axis. This is also consistent with the expression of known markers of the medial (e.g. *Bmp4*, *Isl1*, and *Tlx1*) and lateral (*Fgf8*) portions of the mandible [10,12,72,73], which are enriched in the bottom and the top halves of the UMAP respectively (Fig. S2E-H, Fig. S4K).

**Figure 3.**
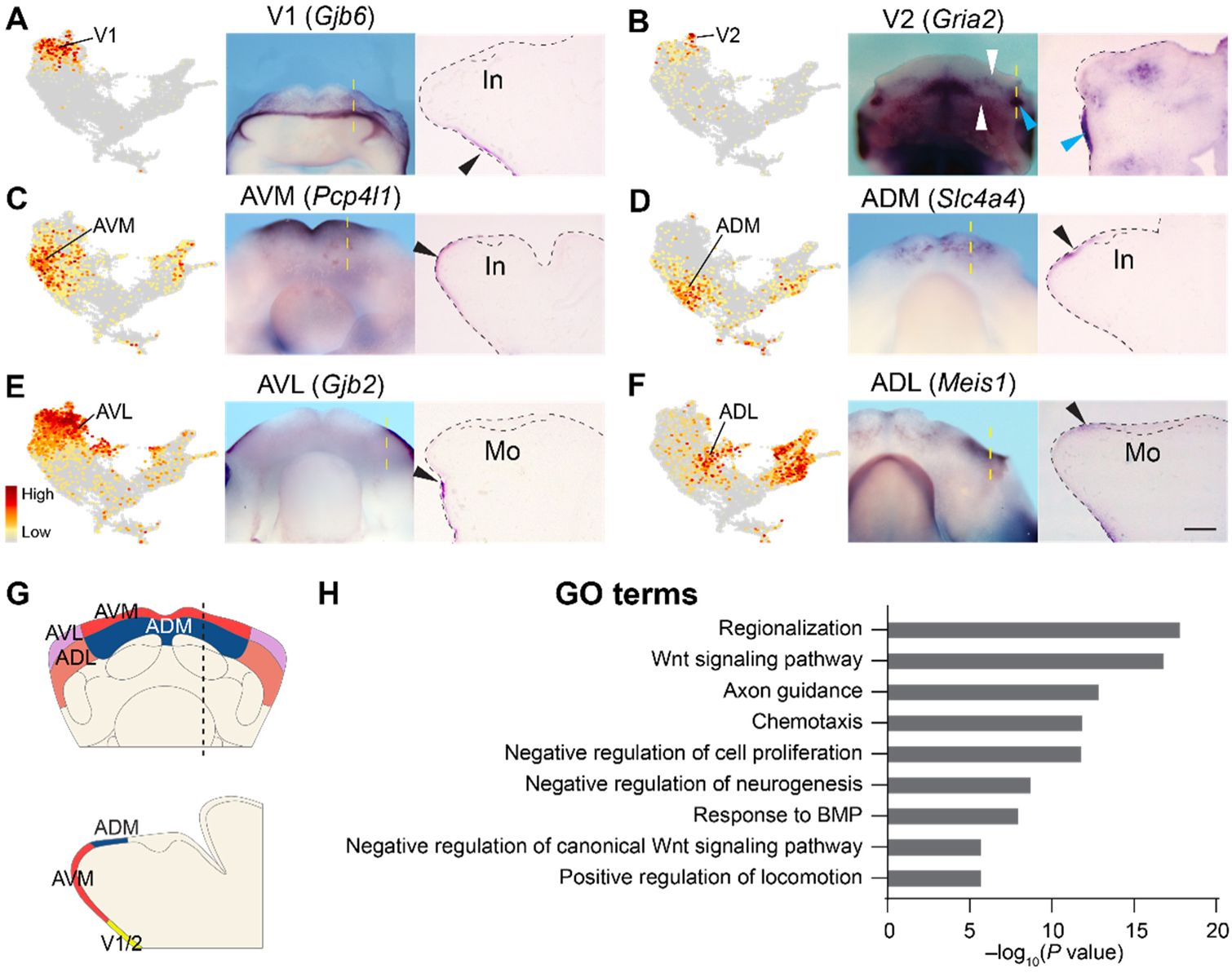
Cell clusters in the anterior mandibular epithelium represent subdomains of the region. (**A-F**) *Gjb6* (A), *Gria2* (B), *Pcp4l1* (C), *Slc4a4* (D), *Gjb2* (E), and *Meis1* (F) are marker genes for the ventral and anterior clusters V1 (ventral 1), V2 (ventral 2), AVM (anteroventral-medial), ADM (anterodorsal-medial), AVL (anteroventral-lateral), and ADL (anterodorsal-lateral) respectively. Their expression are shown in feature plots (left panels) and detected in E12.0 mandibles by whole mount RNA *in situ* hybridization (middle panels), viewed ventrally (A,B) or dorsally (C-F). Right panels are representative sagittal sections through the yellow dashed lines; anterior to the left. Black arrows indicate epithelial expression on sections. *Gria2* is detected as puncta (white arrowheads) on the ventral surface and in a lateral patch (cyan arrowheads). Black dashed lines outline the mandible and the dental epithelium. (**G**) Schematic drawings of the mandible in dorsal view and sagittal section summarizing the distribution of clusters in the anterior and ventral mandibular epithelium. (**H**) Bar graph showing enriched GO terms in the anterior mandibular clusters. *P*-values are generated by Metascape using cumulative hypergeometric distributions. In, incisor; Mo, Molar. Scale bar in (F) represents 450 μm in whole mount images of (A,B); 280 μm in whole mount images of (C-F); and 100 μm in cross-section images of (A-F).

To probe for the functional importance of genes expressed in these clusters, we performed functional enrichment analysis [55]. As the outputs from individual clusters were comparable, the anterior epithelial populations collectively perform similar functions, which are summarized in Figure 3H. We found that there is an overrepresentation of genes related to WNT signaling, and that the anterior mandibular epithelium is a major source of WNT ligands, expressing *Wnt3*, *4*, *5a*, *6*, *7a*, *7b*, *9b*, *10a*, and *10b* (Table S2). Several WNT inhibitors, *Axin2*, *Znrf3*, *Kremen2*, *Nkd1*, and *Sostdc1*, are also upregulated, likely as a part of the negative feedback loop downstream of WNT signaling [74–77]. In parallel, anterior epithelial cells are capable of mediating and modulating signals from BMP and TGFβ, as they express a group of genes that include *Msx1*/*2*, *Nbl1*, and *Htra1* [78–80]. Finally, another category of genes in the anterior mandibular epithelium encode regulators of chemotaxis, such as *Ephb1/2*, *Robo2*, and *Sema3a*, and they may contribute to axon guidance [81–84] and/or the migration of other cell types in the anterior developing mandible.

### scRNA-seq identifies several novel markers for different tooth-related populations

We next focused on clusters with dental signatures. Examining genes highly expressed in the cluster IK revealed known markers (e.g. *Shh*, *Dkk4*, and *Fgf20*) of the tooth signaling center, the initiation knot (Fig. S4A) [23]. We have also identified and validated several novel IK markers including *Gad1*, *Sp5,* and *Proser2* (Fig. 4A, Fig. S4C,D), and as expected IK is enriched with components of several signaling pathways that promote cell proliferation and tooth development (Fig. 4F).

**Figure 4.**
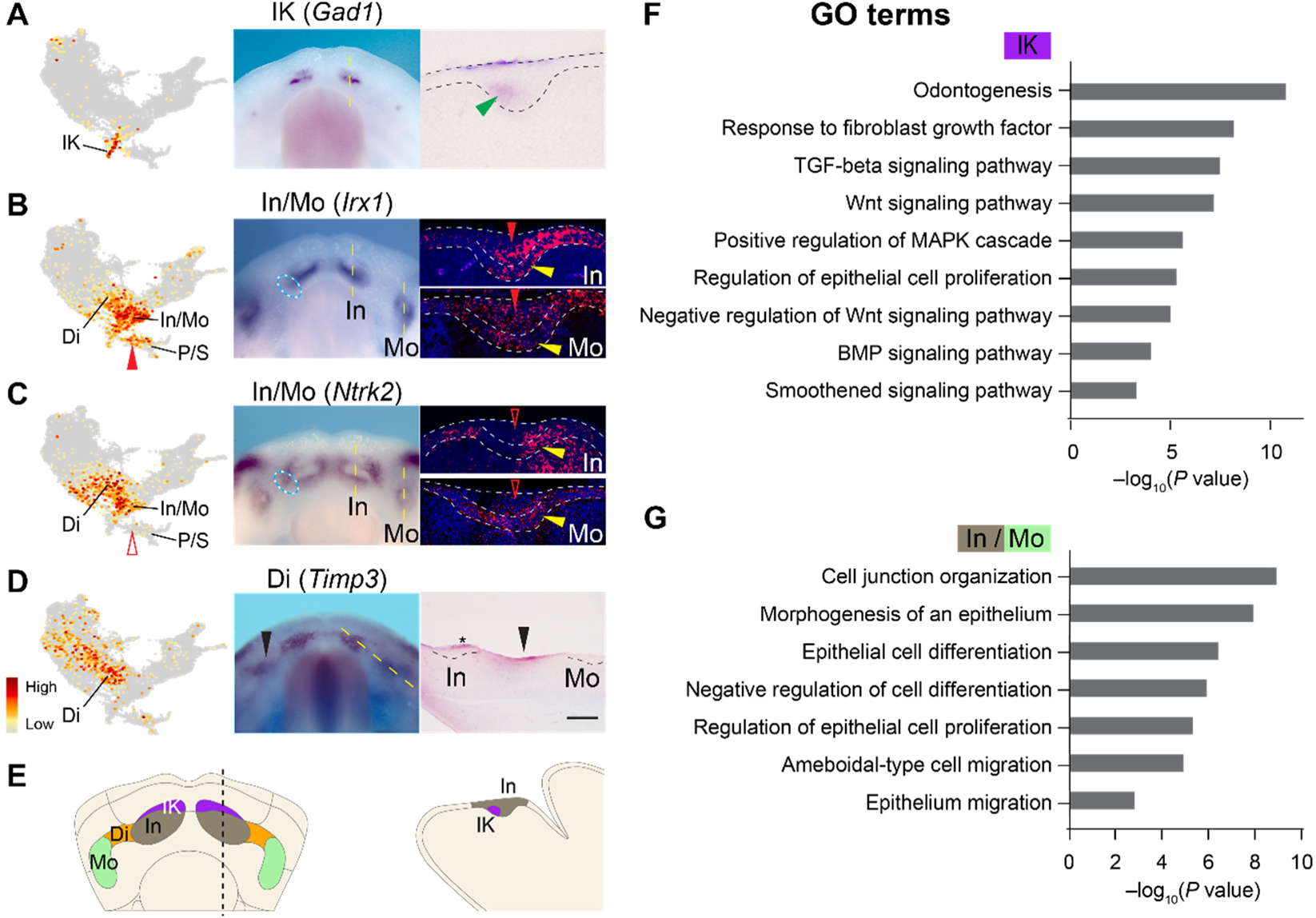
Distinct cell populations reside in the dental lamina and express genes important for epithelial morphogenesis. (**A-D**) Feature plots (left panels) and RNA *in situ* hybridization on E12.0 mandibles (middle panels, dorsal views) for markers enriched in dental-related clusters. Sagittal sections (right panels) show representative *in situ* (A,D) or RNAscope (B,C) staining at the levels indicated by the yellow dashed lines; anterior to the left. Black and white dashed lines outline the dental epithelium. (A) IK marker *Gad1* labels the initiation knot (green arrowhead). (B,C) *Irx1* and *Ntrk2* are enriched in clusters In and Mo, corresponding to their expression in the non-IK basal layer of the incisor and the molar (yellow arrowheads); and in parts of cluster Di, corresponding to the expression spreading into the diastema (cyan ovals). Solid and open red arrowheads indicate strong or weak expression in the suprabasal cells, contained within the P/S (periderm and suprabasal) cluster. (D) Di marker *Timp3* labels the diastema region (black arrowheads) and periderm cells over the incisor (asterisk). (**E**) Summary of results from *in situ* hybridization mapping of dental-related clusters. (**F,G**) Bar graph showing enriched GO terms in the IK and In/Mo clusters. *P*-values are generated by Metascape using cumulative hypergeometric distributions. Scale bar in (D) represents 280 μm in all whole mount images, 50 μm in cross-section images in (A-C), and 100 μm in cross-section image in (D).

Juxtaposing with the cluster IK on the UMAP are clusters In and Mo, which have similar transcriptional profiles (Fig. 2C). Among the genes that are highly expressed in both In and Mo are known dental markers (e.g. *Pitx2, Irx1*, and *Fst*) [17,18,85], as well as newly identified genes (e.g. *Ntrk2*, *Dsc3*, *Enc1*, and *Osbpl6*), whose expression we verified in the developing teeth (Fig. 4B,C, Fig. S4E-H). To further distinguish the identities between the two clusters, we examined genes that are uniquely expressed in cluster In. This identified *Ednrb* and *Ddit4l*, which are localized to the incisor based on *in situ* hybridization (Fig. S4I,J). Cluster Mo must then represent the molar epithelium, as supported by its expression of the molar marker, *Fgf8* (Fig. S4K). Notably, many dental markers are expressed beyond just the In and Mo clusters on the UMAP. For example, *Dsc3* and *Enc1* are additionally expressed in cluster IK and their transcripts are present throughout the entire incisor bud at E12.0 (Fig. S4F,G). In contrast, expression of *Irx1* and *Ntrk2* is considerably lower in the IK cluster and cells expressing these genes correspondingly occupy the non-IK portion of the incisor bud, as assessed using RNAscope *in situ* hybridization (Fig. 4B,C). This is consistent with the expression of *Sox2*, which was shown to label the posterior incisor bud at E12.0 [63] and is differentially expressed in the In/Mo clusters but not the IK cluster (Fig. S4B). In addition, we noticed that many In/Mo markers are also detected at high levels in cluster P/S, and this corresponds to their expression in both the basal and suprabasal cells of the developing tooth bud (e.g. *Irx1* and *Dsc3* in Fig. 4B and Fig. S4F). In contrast, In/Mo markers with minimal presence in cluster P/S are localized primarily to just the dental basal layer (e.g. *Ntrk2* in Fig. 4C). Therefore, cells in clusters In and Mo most likely represent the dental basal cells, while the suprabasal cells are transcriptionally closer to the peridermal cells in cluster P/S, of which we will describe in a later section. At the functional level, many of the genes in the In/Mo clusters are involved in regulating cell proliferation and differentiation (Fig. 4G), and this is consistent with the general role of the epithelial basal layer, where progenitor cells divide and give rise to more differentiated suprabasal cells [28,86]. Interestingly, regulators of cell protrusions and adhesions, such as *Slitrk6*, *Wasf1*, *Dock5*, *Flrt3*, and *Ednrb* are specifically upregulated in In/Mo cells (Table S2) and they may control the process of basal cell delamination into the suprabasal layer [87–91].

Lastly, we examined the expression of Di markers *Timp3* and *Prss23*, and found that they mark the space between the incisor and molar buds (Fig. 4D and Fig. S4L), thus mapping cluster Di to the diastema. In support of this, several In/Mo genes (e.g. *Irx1* and *Ntrk2*) are additionally enriched in parts of the cluster Di on the UMAP, and their whole mount *in situ* staining displayed corresponding expression that extends from the tooth bud into the diastema region (cyan ovals in Fig. 4B,C). The assignment of tooth-related clusters is summarized in Fig. 4E.

### Suprabasal populations are diverse and exhibit transcriptional features for strong cell-cell adhesion and cell movement

Our analysis so far indicates that cluster P/S contains both peridermal and suprabasal cells. To verify this, we examined the expression of P/S-specific markers, *Ppl*, *Marveld2*, and *Cpm*, and found that they label both suprabasal and peridermal cells in the tooth bud, but are excluded from the basal layer (Fig. 5A-D). This result thus confirms that the dental suprabasal cells are grouped within the P/S cluster. Functional enrichment analysis revealed that P/S cells are characterized by tight junction and desmosome genes (e.g. *Cldn4*, *Cldn7*, *Jup*, *Ppl*) (Fig. 5E), which are often associated with differentiating cells in stratified epithelia [92]. At the level of signaling regulation, *Notch1* and *Notch3* are expressed in the P/S, while their ligand genes *Jag1* and *Jag2* are additionally expressed in the underlying basal layer (Fig. S5A). This is therefore in line with reports showing that the generation of periderm and suprabasal cells depends on Notch signaling [93,94]. Finally, P/S cells are enriched with genes encoding factors that organize actin cytoskeletons and promote cell motility (e.g. *Limk2*, *Pdlim5*, *Csrp1*, *Myl9*, *Rhov*) (Fig. 5E). This would enable suprabasal cells to actively move and generate tissue contraction forces that are needed for tooth epithelial invagination as previously discovered [95].

**Figure 5.**
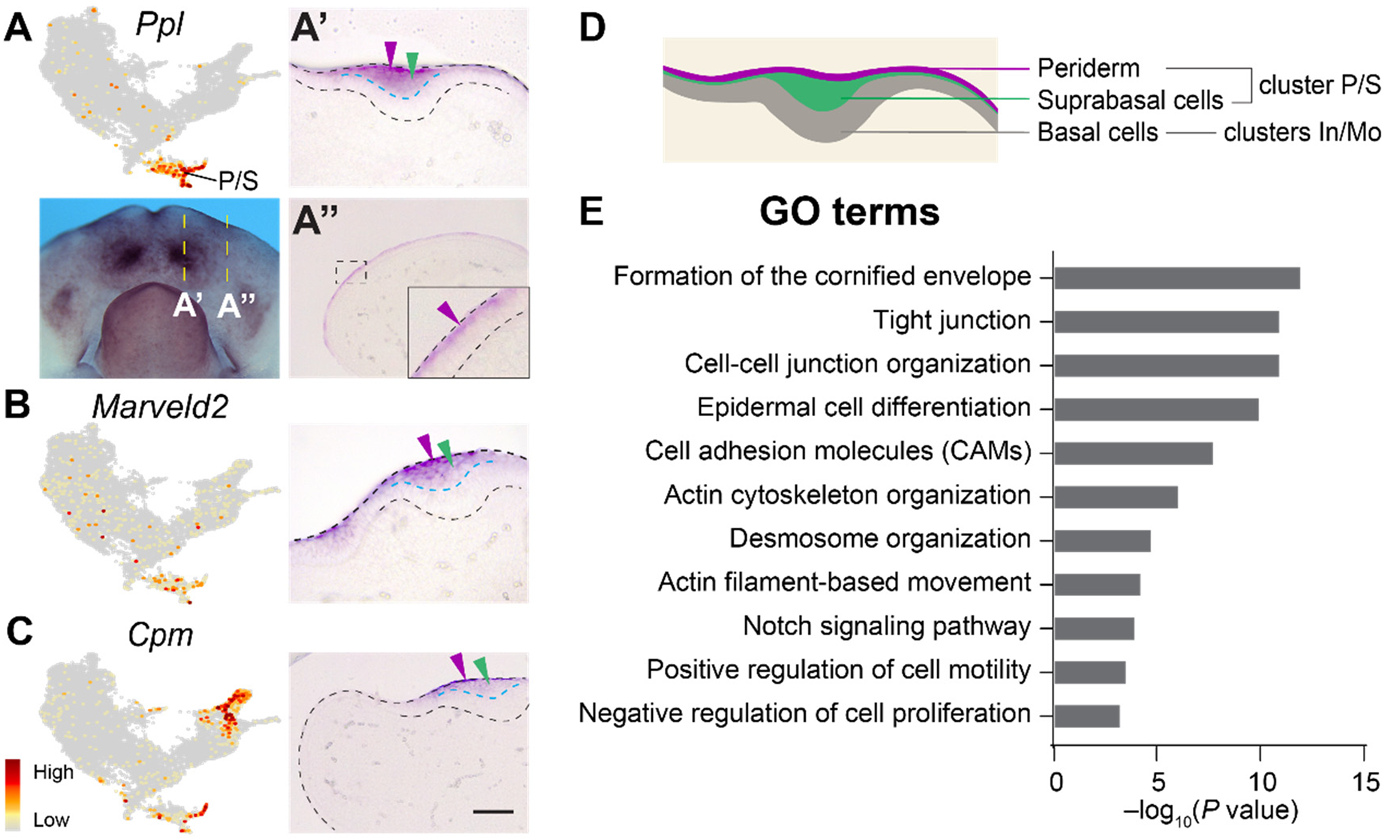
Periderm and suprabasal (P/S) cells are transcriptionally distinct from basal cells and primed for cell-cell adhesion. (**A-C**) Feature plots show *Ppl* (A), *Marveld2* (B), and *Cpm* (C) as P/S markers. *In situ* hybridization on sagittal sections (anterior to the left) show their expression in both the periderm (purple arrowheads) and the suprabasal cells (green arrowheads) of the dental placode, but not in the basal layer (below the cyan dashed lines). (A’,A”) are sections through the yellow dashed lines. Black dashed lines outline the mandible and the incisor bud. (**D**) Schematic of a dental placode showing different populations and their corresponding clusters. (**E**) Bar graph showing enriched GO terms in cluster P/S. *P*-values are generated by Metascape using cumulative hypergeometric distributions. Scale bar in (C) represents 280 μm in (A); 50 μm in (A’,B,C); 100 μm in (A’’); and 25 μm in the inset of (A’’).

We next questioned whether there is a transcriptionally distinct periderm population that is discrete from the suprabasal cells. To dissect the heterogeneity among P/S cells, we performed sub-clustering and identified 4 sub-populations (P/S1-4) (Fig. S5B, Table S3). Intriguingly, P/S1-3 mirror the oral-aboral patterning we have observed in the rest of the mandibular epithelium. They express the same positional markers and respectively cover the aboral (P/S1, Fig. S5C,D), anterior oral and dental (P/S2, Figs. S4F,L, S5E), as well as the posterior oral (P/S3, Fig. S5F-H) epithelium, which we describe next. On the other hand, P/S4 is comprised of differentiated peridermal cells, which express novel markers *Tagln* and *Acta2*, and are only observed on the apical surface of the mandibular epithelium (Fig. S5I,J). Therefore, P/S cells are diverse, and their transcriptional variations reflect differences in their differentiation progression and/or localization.

### E12.0 tongue epithelium is a transcriptionally distinct population that includes precursor cells of taste bud primordia

The rest of the UMAP contains epithelial cells posterior to the dental tissues, encompassing clusters PM, PL, SG, and T. Cluster PM expresses markers *Dmrt2*, *Wisp1*, and *Rprm*, which are mapped to the space between the tooth and the tongue (Fig. 6A,F, Figs. S5H, S6A,B). PM cells also express a suite of genes (e.g. *Meis1*, *Sox2*, and *Smarca2*) known to maintain the progenitor state of cells [96–98] (Fig. 6G) and could function here to limit epithelial differentiation and stratification. Cluster SG (marked by *Hpca*, *Col14a1*, *Col9a1*, and *Hpgd*) represents the invaginating salivary gland bud [99] (Fig. 6B, Fig. S6D-F), while cluster PL (marked by *Fndc1*, *Ntn1, Pax9,* and *Foxc1*) includes junctional cells lateral to the tongue that connect the salivary gland bud to the mandibular and tongue epithelium (Fig. 6C, Fig. S6G-I). Functional enrichment analysis showed that cells in SG/PL are enriched with regulatory genes shared among glandular and ductal organs (e.g. *Six1*, *Six2*, and *Foxc1*) (Fig. 6H), highlighting a common mechanism for branching morphogenesis [100,101]. In parallel, the upregulation of genes associated with FGF signaling and programmed cell death is concordant with the developmental process of salivary gland lumen formation [102].

**Figure 6.**
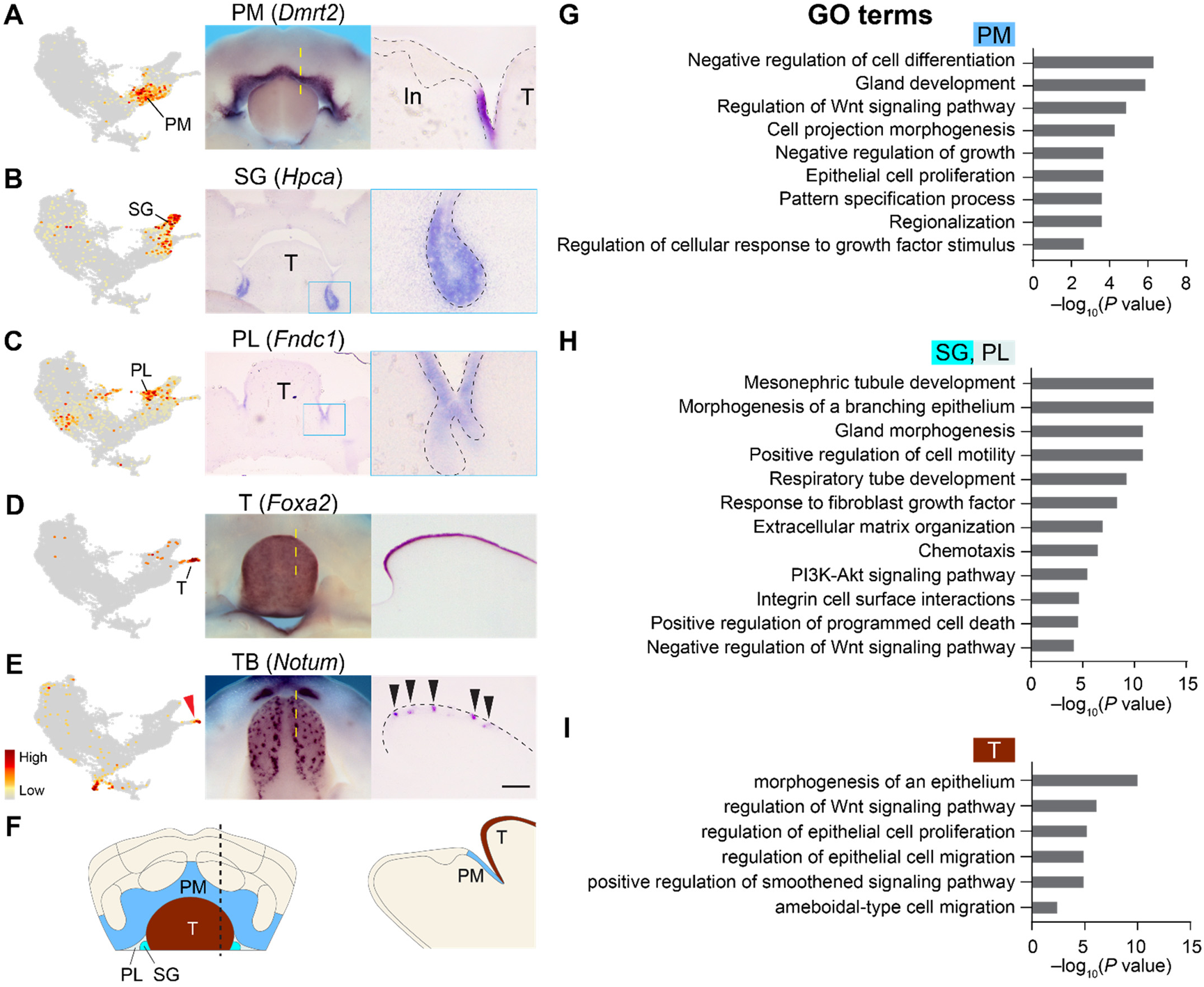
Posterior mandibular clusters include cells in the salivary gland and the taste bud primordia. (**A-D**) Left panels show feature plots of indicated marker genes for the posterior epithelial clusters PM (posterior-medial), SG (salivary gland), PL (posterior-lateral), and T (tongue). Middle panels show *in situ* hybridization staining on E12.0 whole mount mandibles (A,D, dorsal views) or frontal sections (B,C). Right panels are sagittal sections (anterior to the left) through the yellow dashed lines in (A,D) or zoomed in images in (B,C). (**E**) The tip of the tongue cluster (red arrowhead) contains taste bud (TB) precursor cells, here detected by *in situ* hybridization of *Notum* on a E12.5 mandible and its sagittal section. Black arrowheads mark taste bud placodes. (**F**) Summary diagrams of the posterior mandibular epithelial populations. (**G-I**) Bar graphs showing enriched GO terms in clusters PM, SG/PL, and T. *P*-values are generated by Metascape using cumulative hypergeometric distributions. T, tongue; In, incisor. Scale bar in (E) represents 280 μm in whole mount images in (A,D,E); 200 μm in cross-section images in middle panels of (B,C), 50 μm in right panels of (A-C), and 100 μm in cross-section images in (D,E).

The tongue epithelial cells are distributed in cluster T and are labelled by *Foxa1* and *Foxa2* (Fig. 6D, Fig. S6C). Curiously, *Shh*, a known marker of the forming taste buds [103], is specifically restricted to the distal tip of cluster T on the UMAP (Fig. S7B). We therefore performed sub-clustering and identified additional markers for this subpopulation (Table S4), which is likely composed of precursor cells of the taste bud primordia (Fig. S7A). These markers, which include *Notum*, *Dsc3*, *Sp6*, and *Prss23*, are expressed at a low level on the tongue surface at E12.0, when the taste bud primordia are just beginning to form [104]. They then become more discernable at E12.5 (Fig. 6E, Fig. S7C-E). Interestingly, taste bud and dental placodes co-express several markers (Fig. S7F), indicative of similar developmental processes and signaling regulations (Figs. 4F,G and 6I). On the contrary, salivary glands share fewer markers with either the taste bud or the tooth at E12.0 (Fig. S7F), instead employing other mechanisms to form tubules and branched structures.

### Mandibular epithelium undergoes progressive regionalization along the oral-aboral axis

Our analyses so far established the spatial pattern of different mandibular epithelial populations at E12.0. The identification of several region-specific markers provided an opportunity to study how the tooth and its neighboring epithelium are patterned along the oral-aboral axis over time, which remains not well understood. At E12.0, both *Irx1* and *Pitx2* are robust markers of the developing teeth (Figs. 1F and 4B) [67]. Based on the UMAP and our *in situ* mapping of individual genes, the *Irx1*+ and *Pitx2*+ dental domain should in theory be bounded anteriorly by cells in the ADM/AVM/ADL/AVL clusters that collectively express *Cxcl14* and *Tfap2b* and posteriorly by *Dmrt2*-expressing PM cells (Fig. 2). To simultaneously visualize these markers and compartments, we performed RNAscope *in situ* hybridization on sagittal sections at the level of the incisor bud. Using the combination of *Cxcl14*/*Irx1*/*Dmrt2* and *Tfap2b*/*Pitx2*/*Shh*, where *Shh* is a marker for both the initiation knot and taste buds, we were able to concretely show that at E12.0 the mandibular epithelium is divided into 3 main zones: the zone anterior to the dental epithelium (zone A), the dental zone (zone D), and the zone posterior to the tooth (zone P) (Fig. 7F,L,U). At the junctions between these zones, cells co-express markers from the neighboring regions. For example, *Cxcl14+*/*Irx1+* and *Irx1+*/*Dmrt2+* cells respectively span the A/D and D/P interzone boundaries, which at this stage consist of 1-3 cells (Fig. 7M,N). On the other hand, as *Pitx2* expression is broader than *Irx1*, the A/D boundary labelled by *Tfap2b*/*Pitx2* is also comparably wider (Fig. 7O).

**Figure 7.**
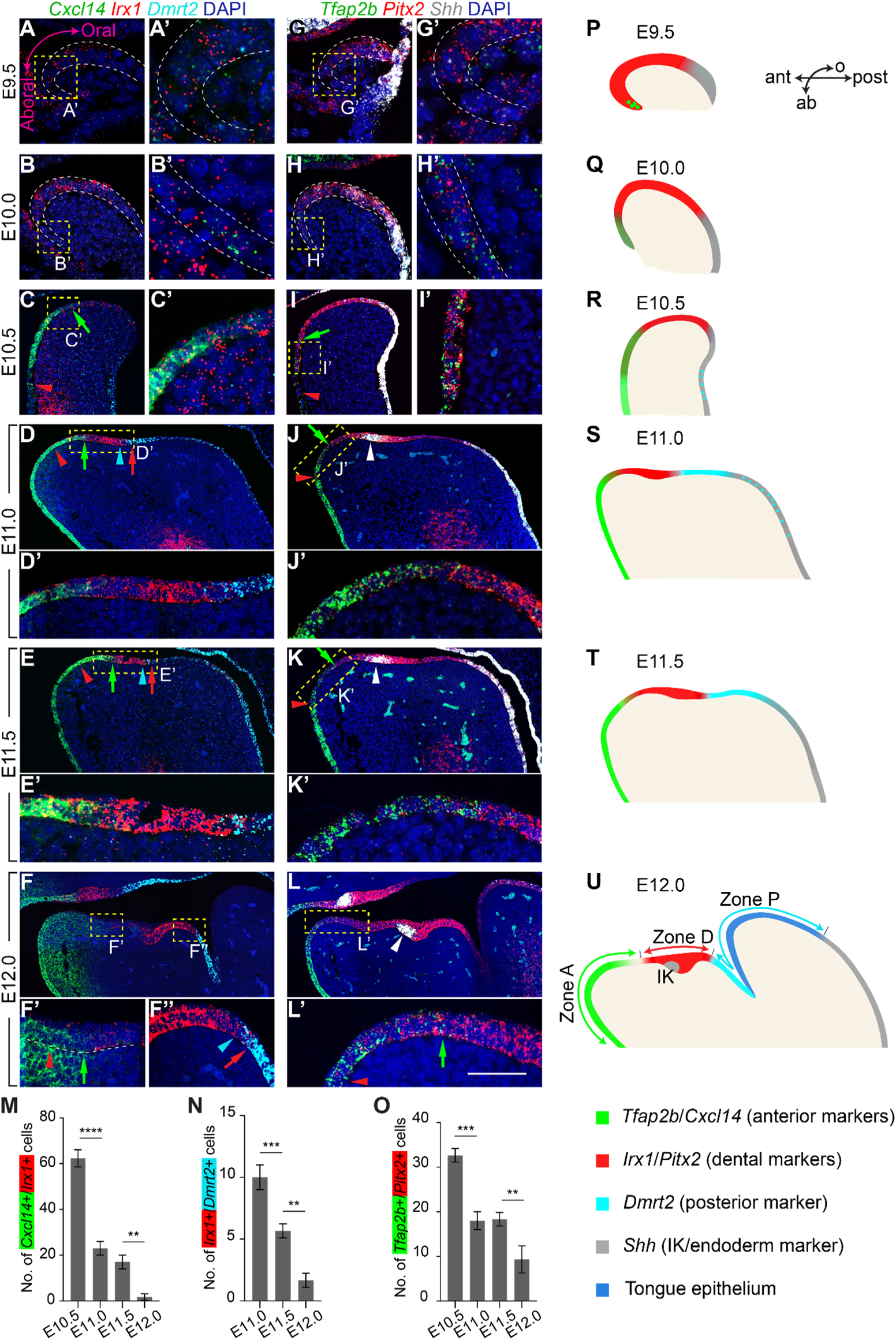
The mandibular epithelium undergoes progressive transcriptional regionalization along the oral-aboral axis from E9.5 to E12.0. (**A-L**) RNAscope analysis showing spatiotemporal expression changes of markers for the anterior mandibular epithelium *(Cxcl14* and *Tfap2b*, green), the dental epithelium (*Irx1* and *Pitx2*, red), the posterior epithelium (*Dmrt2*, cyan), and the oropharyngeal endoderm (*Shh*, white) along the oral-aboral axis from E9.5 to E12.0. *Shh* also labels the initiation knot (white arrowheads). Representative sagittal sections through the presumptive incisor region (A-D’,G-J’) or the incisor bud (E-F”,K-L’) are shown; anterior to the left. (A’-L’) are enlargements of yellow dashed boxes in (A-L). White dashed lines outline the ectodermal layer. Colored (red, green, and cyan) arrowheads and arrows respectively mark the anterior and posterior limit of cells labelled by the same color, and they denote the spread of boundary cells that have overlapping expression of markers from the adjoining regions. (**M-O**) Quantification of the boundary cell numbers at indicated timepoints (n=3). (**P-U**) Schematics summarizing the expression pattern of regional markers and the establishment of different epithelial zones along the oral (o)-aboral (ab) and anterior (ant)-posterior (post) axes of the mandible from E9.5 to E12.0. Zone A, anterior zone; Zone D, dental zone; Zone P, posterior zone; IK, initiation knot. Scale bar in (L’) represents 60 μm in (A,G,C’-E’,I’-K’); 100 μm in (B,H,F’,F”,L’); 200 μm in (C-F,I-L); and 20 μm in (A’,B’,G’,H’). Quantitative data are presented as mean ± SD. *P*-values are determined using one-way ANOVA and Tukey’s HSD test. **p<0.01; ***p<0.001; **** p < 0.0001.

To examine how the spatial pattern of the three zones, as defined by their markers, may change over time during mandibular development, we analyzed the expression of the same set of genes in younger embryos between E9.5 and E11.5. It should be noted that *Shh* is used as a marker for the oropharyngeal endoderm at these earlier stages [13,105,106] and to contrast the expression of other markers in the ectoderm. Lineage tracing with *Shh^CreER^;R26^mT/mG^* confirmed that cells labelled between E8.5 and E10.5 are restricted to the endoderm and do not give rise to epithelium beyond the posterior third of the tongue (Fig. S8A), thus consistent with an earlier study using *Sox17-2A-iCre* [9]. Interestingly, at E9.5 when the mandibular epithelium is just a monolayer, *Irx1* and *Pitx2* are present in the majority of ectodermal cells and border the *Shh*+ endoderm (Fig. 7A,G,P). On the contrary, markers for zones A and P are barely detectable at this stage. At E10.0, a small group of *Irx1*+/*Pitx2*+ cells at the ventral mandible next to the developing heart begin to also express zone A markers, *Cxcl14* and *Tfap2b* (Fig. 7B,H,Q). Their expression then becomes noticeably expanded between E10.0 and E10.5, coinciding with the rostrocaudal elongation of the mandible (Fig. 7C,I,R). *Irx1* and *Pitx2* expression are now located at the oral surface, but the *Cxcl14+*/*Irx1+* and *Tfap2b+*/*Pitx2+* boundaries remain relatively diffused across about 62 and 33 cells respectively (Fig. 7M,O). Posteriorly, *Irx1* and *Pitx2* continue to adjoin the endoderm that is labeled by *Shh* at E10.5 (Fig. 7C,I,R). By E11.0 *Dmrt2*+ cells finally emerge in the ectoderm between the dental lamina and the oropharyngeal endoderm, encompassing the region that would form zone P and the anterior part of the tongue (Fig. 7D,S, Fig. S8B,C). We made a similar observation using *Zcchc5* as a marker for the posterior oral epithelium (Fig. S8D,E). The expression of zone P markers at E11.0 thus delineates *Irx1* in the dental lamina posteriorly, and a P/D boundary is established. Both A/D and P/D boundaries continue to narrow at subsequent stages (Fig. 7D-F,J-O), such that from E10.5 to E12.0 the interzone boundaries are progressively sharpened, just as the 3 zones become increasingly defined (Fig. 7M-U).

Comparing RNAscope results from E10.5 to E12.0, we noticed that the expression of zone A and zone D markers at the A/D boundary are considerably weaker at E12.0 than at earlier stages. As this is where ADM cells are located, we reasoned that the ADM fate is newly specified around E12.0 and ADM cells would be absent at earlier timepoints. To test this, we compared our E12.0 scRNA-seq data with a published E10.5 dataset [13]. This revealed that most zone A cells at E10.5 are transcriptionally similar to V1, AVM, and AVL, and only a small portion of them are clustered with ADM and ADL (Fig. S8F,G), thus supporting the idea that ADM and ADL are later stage populations that are specified in between the aboral epithelium and the forming tooth bud. The same result was obtained by examining the expression of *Rxfp1*, which is a marker for both the ADM and zone P at E12.0 (Fig. 2C). However, while *Rxfp1* is expressed in zone P labeled by *Dmrt2* and *Zcchc5* beginning at E11.0, *Rxfp1* expression in the anterior-dorsal region only becomes apparent at E12.0, indicating the specification of ADM cells at that stage (Fig. S8C,E). We observed a similar trend of sequential formation for the PM cluster, as zone P markers are only expressed after the formation of zones A and D (Fig. 7), and the proportion of PM cells at E10.5 is considerably lower than at E12.0 (Fig. S8F,G). Together, these results demonstrated that mandibular patterning along the oral-aboral axis is a dynamic regionalization process, and the initial broad expression of dental markers are progressively delimited anteriorly and posteriorly by newly specified epithelial populations between E9.5 and E12.0. Applying Slingshot to infer pseudotime lineage trajectories on the E10.5 scRNA-seq data showed that *Pitx2*+ and *Irx1*+ cells could give rise to *Cxcl14*+ and *Tfap2b*+ cells (Fig. S8H), supporting the idea that aboral cells arise as they gradually shift from a dental-like identity to the zone A fate.

### Region-specific transcription factors underlie the transcriptional differences between epithelial populations

To gain insight into the gene regulatory network (regulon) that defines the different mandibular epithelial populations, we next applied the SCENIC pipeline [59] on our E12.0 dataset to computationally infer key transcription factors and their downstream target genes based on co-expression and enrichment of *cis*-regulatory motifs. This yielded region- and cluster-specific regulons (Fig. 8, Table S5), which include many of the marker genes we described earlier. For instance, *Gata3*, *Hoxc13*, *Lef1*, and *Trps1* all encode transcription factors, and they are differentially expressed in the V1/V2/AVM/AVL cells of the anterior/aboral epithelium (Fig. 8A,B). Many markers of these clusters, such as *Gjb2* and *Skap2* (Fig. 8A), correspondingly contain binding motifs for these transcription factors. The detection of a LEF1 regulon echoes our findings that active Wnt signaling is a key feature of the cell state in the anterior/aboral epithelium at E12.0 (Fig. 3H). Also notably, TRPS1 is an atypical GATA transcription factor that can recognize the GATA-binding sequence but functions as a transcriptional repressor [107]. As TRPS1 and GATA3 regulons share several targets (Fig. 8A) and their mutations can affect mandibular patterning and tooth formation [108,109], they may modulate the expression of a gene set to help specify the aboral mandible. In contrast, *Nkx2-3* is expressed in the oral epithelium (Fig. 8A,C). It targets several other transcription factors, such as *Irx1*, *Pitx2*, *Foxp2*, and *Sox21*, all of which have their own regulons (Fig. 8A). IRX1 and PITX2 are the predicted main transcription factors in the dental epithelium (Fig. 8A,D, Fig. S9A), while FOXP2 and SOX21, together with SOX2 and TCF7L2, target genes in the more posterior oral epithelium (Fig. 8A,G, Fig. S9D).

**Figure 8.**
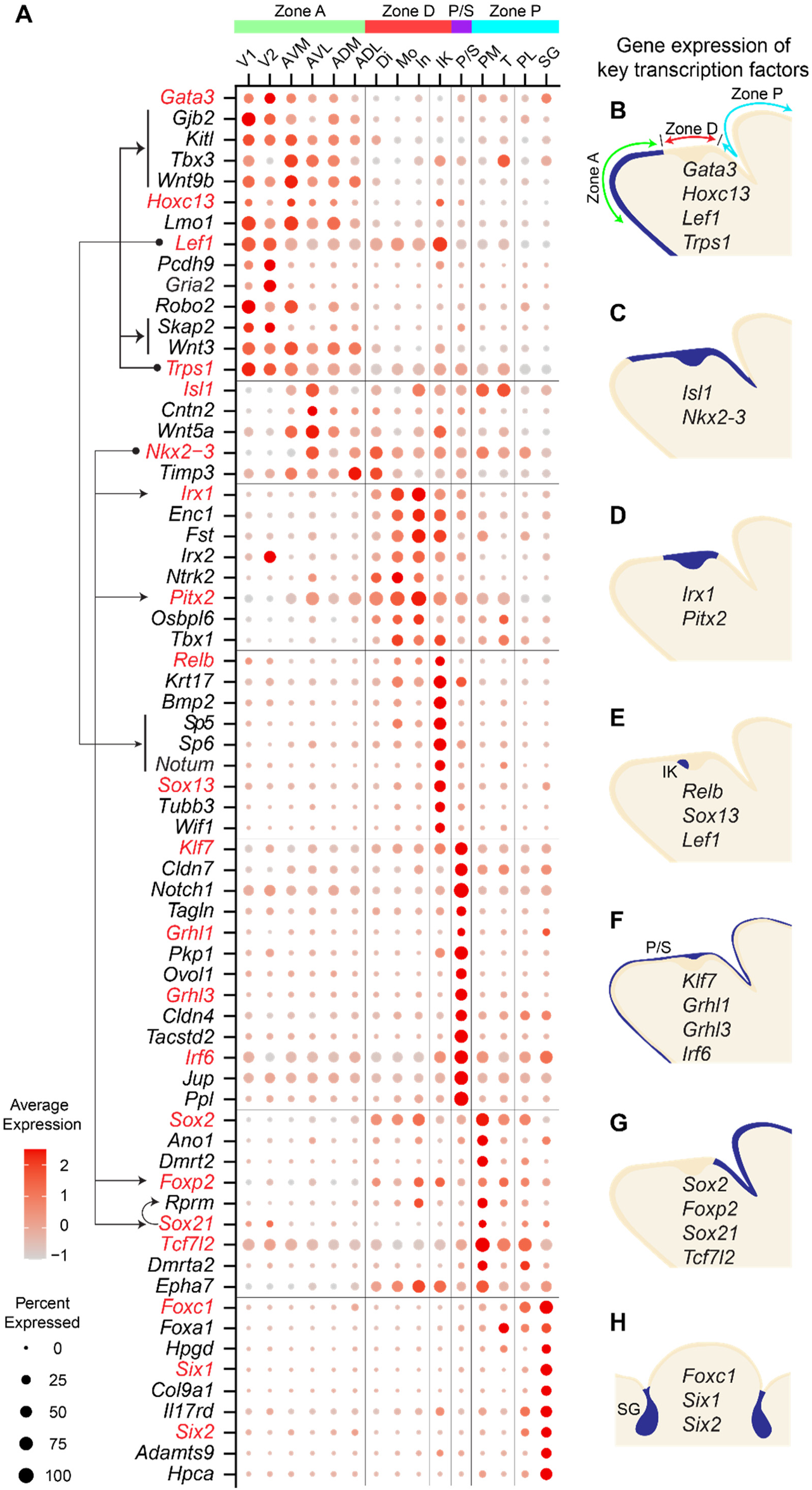
SCENIC analysis reveals population-specific transcription factors and downstream targets. (**A**) Dot plot showing the expression of genes encoding key transcription factors (labelled in red) and examples of downstream targets (listed below each transcription factor) in each cluster and epithelial regions, as identified by the SCENIC pipeline. Genes targeted by multiple transcription factors are connected by arrows. (**B-H**) Schematics of sagittal sections through the incisor (B-G) or of a frontal section through the submandibular glands (H) depicting the RNA expression of key transcription factors in specific epithelial regions; anterior to the left. Zone A, anterior zone; Zone D, dental zone; Zone P, posterior zone; IK, initiation knot; P/S, periderm and suprabasal cells; SG, salivary gland.

Our SCENIC analysis also uncovered regulons by RELB, LEF1, and SOX13 in the initiation knot (Fig. 8A,E and Fig. S9B). RELB is a transcription factor within the NF-κB pathway and has been implicated to function downstream of the Eda/Edar signaling to control the number of initiation knot cells [23]. Concurrently, WNT activity has been proposed to promote initiation knot formation [24] and our result here supports this view. In the periderm, we have identified regulons associated with the transcription factors GRHL1, GRHL3, and IRF6 (Fig. 8A,F), which all play critical roles in driving periderm differentiation [69–71]. This is in line with the presence of several tight junction and desmosome components as their transcriptional targets (Fig. 8A and Fig. S9C), many of which also contain motifs for KLF7, a Krüppel-like transcription factor. KLF7 therefore represents a potential novel regulator of the periderm population. Lastly, we have identified FOXC1, SIX1, and SIX2 as the key transcription factors for the developing salivary gland (Fig. 8A,H, Fig. S9E). Importantly, these factors are both required for the development of several ectodermally-derived glandular organs, including salivary and lacrimal glands [100,101], and also critical for inducing mouse embryonic stem cell-derived oral ectoderm into gland organoids [110,111]. This thus suggests a common regulatory program controlled by these factors to direct gland morphogenesis. Together, our analysis here revealed multiple gene regulatory networks in the mandibular epithelium and connected their spatial expression to the transcriptional control of distinct epithelial populations.

### NTRK2 promotes epithelial invagination during early tooth development

As our E12.0 scRNA-seq data successfully revealed genes enriched in different oral epithelial appendages, it provides a useful platform to uncover novel regulators of epithelial morphogenesis. As a proof of principle, we focused on the developing tooth and its newly characterized marker *Ntrk2*, which encodes the Neurotrophic receptor tyrosine Kinase 2. NTRK2 is a receptor for brain-derived neurotrophic factor (BDNF) and neurotrophin-4 (NTF4), and its signaling activation regulates proliferation and differentiation in other contexts [112–115]. Because cell proliferation is critical for epithelial stratification during early tooth morphogenesis [28], and the relatively shallow dental lamina begins to grow rapidly in size at E12.0, signaling via NTRK2 may play a role in promoting these processes.

We begin our analysis by first examining the spatiotemporal expression of *Ntrk2* in more detail using RNAscope. Whereas only few *Ntrk2* transcripts were observed in the developing dental epithelium at E11.0 and E11.5, robust expression was detected in both incisors and molars at E12.0 (Fig. S10). In the incisor, *Ntrk2* is especially abundant in the non-IK basal layer and part of the adjacent suprabasal cells, but also in the underlying mesenchyme (Fig. 4C). In the molar, *Ntrk2* is similarly expressed in the basal layer (Fig. 4C). At the same time, *Bdnf* is expressed in the dental epithelium as well as in the anterior part of the mandible (Fig. S10B). *Ntf4* is undetectable by UMAP analysis nor by whole mount *in situ* hybridization at E12.0 (data not shown), indicating that BDNF is the main ligand for NTRK2 signaling during early tooth morphogenesis.

To understand NTRK2 function during early tooth development, we next cultured E11.5 mandible explants in the presence or absence of ANA-12, a selective NTRK2 antagonist [116]. This would block NTRK2 signaling at the onset of its expression and before the rapid growth of a dental placode into a tooth bud. After 3 days of culture, ANA-12-treated samples had significantly smaller tooth buds than controls, both in terms of the depth and the length of the tooth germ (Fig. 9A,B). To examine if cell proliferation was affected following NTRK2 inhibition, we labelled cycling cells by EdU after 2 days of culture. There were significantly fewer EdU+ cells in the suprabasal layer and in the middle portion of the basal layer in ANA-12-treated incisor buds (Fig. 9C,D). Concurrently, as control incisors have already formed the initiation knot, this differentiated region in the anterior control epithelium contained fewer proliferating cells than that of ANA-12-treated samples. Proliferation remained largely unchanged in the mesenchyme and in molar germs (data not shown), indicating that NTRK2 function is tooth type-specific and may control molar morphogenesis through mechanisms independent of proliferation. Together, these results revealed a novel function of NTRK2 to promote cell proliferation and tissue growth in the incisor.

**Figure 9.**
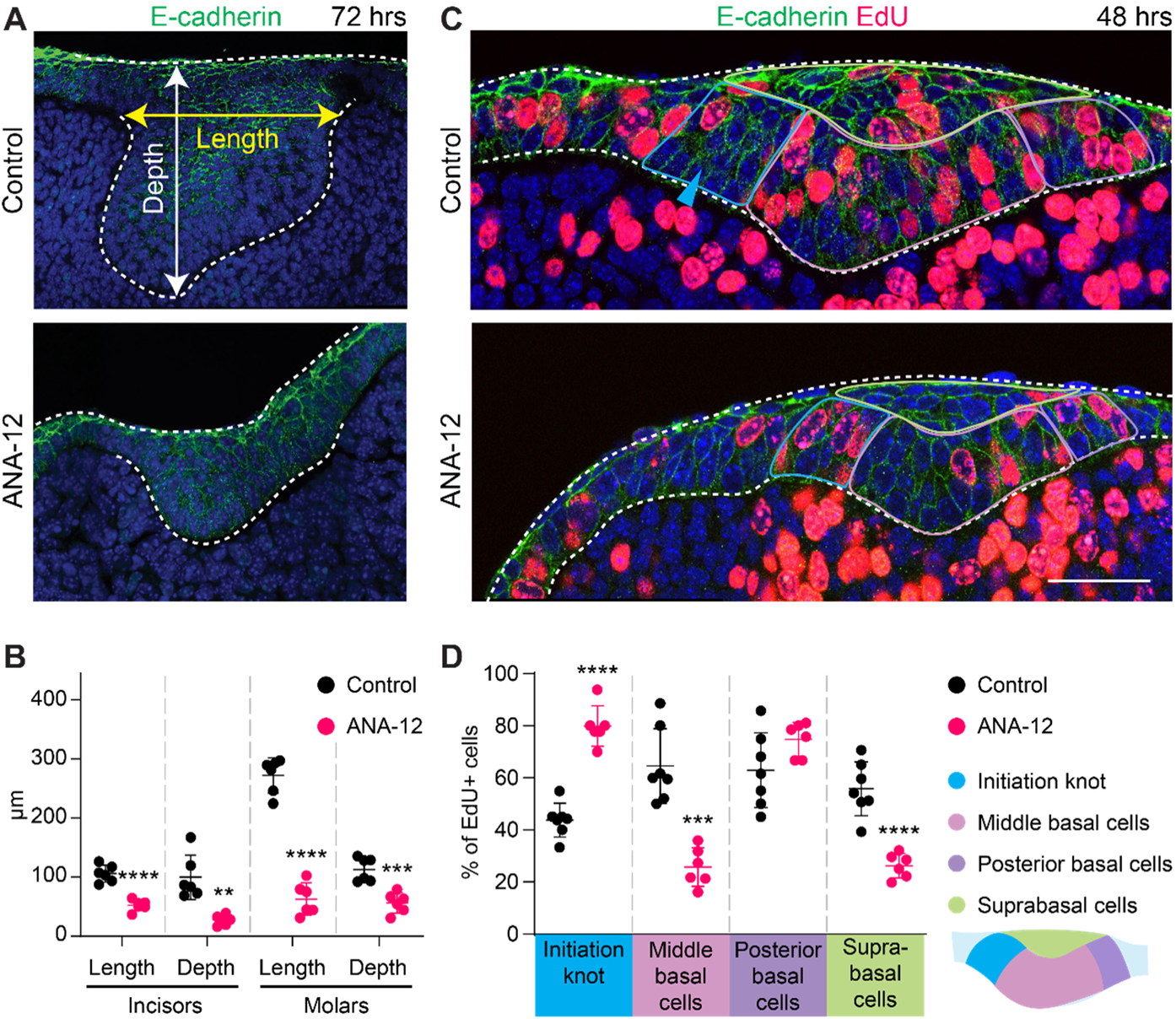
NTRK2 signaling promotes the growth and invagination of the dental epithelium. (**A**) Sagittal sections through the incisor epithelium of E11.5 mandible explants after 72 hours of culture in DMSO control vehicle or the NTRK2 inhibitor ANA-12; anterior to the left. (**B**) Quantification of the length and depth (as shown in A) of the incisor and molar germs treated with DMSO (control) or ANA-12. (n=6). (**C**) EdU staining on sagittal sections through the incisor bud of E11.5 mandible explants treated with DMSO or ANA-12 for 48 hours; anterior to the left. The four sub-regions used for quantifying EdU+ cells are indicated. Cyan arrowhead points at the initiation knot in the control tissue. (**D**) Quantification of EdU+ cells in the four sub-regions of control (n=7) or ANA-12-treated (n=6) incisor epithelium. Dashed lines outline the incisor epithelium, labelled by E-cadherin antibody. Scale bar in (C) represents 80 μm in the control incisor in (A), 50 μm in the ANA-12-treated incisor in (A), and 30 μm in (C). Quantitative data are presented as mean ± SD. *P*-values are determined using Student’s t-tests. **p<0.01; ***p<0.001; ****p<0.0001.

## Discussion

During vertebrate head development, the mandibular ectoderm is remarkable in its ability to give rise to several distinct organs, including the tooth, the salivary gland, the taste bud, and the oral mucosa. This process depends on the correct patterning of the epithelium and specification of different cell types. Using the mouse as a model, our analysis unveiled the transcriptional profiles that define each of the populations comprising the oral epithelium and the different developing structures at E12.0. The resulting atlas not only complements existing knowledge of genes expressed in specific oral structures but also extends previous efforts to profile cells in the developing mandible at other developmental stages using microarray, bulk RNA-seq, or scRNA-seq [33,34,13,35–38]. Our data also highlight the spatial and temporal transitions of gene expression changes that reflect the patterning of the mandibular epithelium along the oral-aboral axis over time. Furthermore, we have identified the gene regulatory networks in different populations and their associated transcription factors that may play key roles in cell fate determination and regulation of cell type-specific functions. Together, this study provides a catalogue of epithelial cell types in the developing mandible and offers a resource for further investigation into the function of specific genes or pathways during epithelial morphogenesis. One such application is our identification of NTRK2 as a promoter of cell proliferation in the invaginating incisor bud.

### Regulation of the mandibular epithelium by WNT

Signaling interactions between the oral epithelium and the underlying mesenchyme as well as within the epithelium itself are critical for the development of oral ectodermal structures [117,118]. Consistent with this, our functional enrichment analysis and gene regulatory network inference showed that many of the differentially expressed genes either encode components of signaling pathways or contain binding motifs for transcription factors responsive to signaling activation (Fig. 8, Fig. S9). For instance, cells in the IK cluster express a multitude of ligands for activating the SHH, BMP/TGFβ, FGF, and WNT pathways (Fig. 4F, Fig. S9B), all of which are crucial for tooth formation [119]. Concurrently, periderm cells express *Notch1-3* and NOTCH target genes, *Hes1*, *Hes5*, and *Hey1* (Fig. S5A), thus supporting a previous finding that active NOTCH signaling maintains the periderm function [93]. Another example is the enrichment of genes downstream of FGF activation in the salivary gland (Fig. 6H), underscoring the importance of FGF signaling during early salivary gland development [5]. We have also identified the anterior and aboral epithelium as a major source of WNT ligands, expressing *Wnt3*, *4*, *5a*, *6*, *7a*, *7b*, *9b*, *10a*, and *10b*. As epithelial WNT9b has been shown to cooperate with R-spondin 2 in the mandibular mesenchyme to promote cell proliferation and survival at E10.5 [120], WNTs at E12.0 may function similarly to enable further mandibular outgrowth. Within the oral epithelium, WNT/β-catenin signaling is a key regulator of tooth formation, as inactivation of the WNT pathway abrogated tooth development [25,121–123] and hyperactivation of WNT signaling led to supernumerary teeth [123–125]. However, WNT responsiveness is not restricted to the dental epithelium, as the expression of WNT target gene *Axin2* and WNT activity reporters both indicate active WNT signaling in the aboral epithelium [65,126], where WNT ligands are highly expressed. This is consistent with our regulon analysis, showing that these cells express *Lef1*, the transcription factor mediating WNT signaling, and are enriched for LEF1 targets. Yet, teeth are normally not formed at the ventral mandible. Therefore, an important question to consider here is how mandibular epithelial cells interpret WNT signals to become different cell types. Paradoxically, while ectodermal deletion of *Tfap2a*/*b* downregulated the expression of multiple WNTs (but not completely lost), it also induced the formation of ectopic incisors at the ventral surface of mutant mandibles [65,66]. The ability to form teeth may thus be modulated by a balance between WNT activity and a yet to be identified inhibitory mechanism downstream of *Tfap2a/b*. We also noticed that several WNT signaling inhibitors, including *Znrf3*, *Kremen2*, *Nkd1*, and *Sostdc1*, are more highly expressed in the aboral epithelium than in the dental epithelium, and could further modulate WNT activities in certain populations.

Another possible mechanism for diversifying WNT responses is by employing different transcription factors. We found that *Tcf7l2* is differentially expressed in the region posterior to the developing tooth and targets many markers in the PM cluster. While LEF1 functions as a transcription activator under high WNT activity, TCF7L2 binds to the same set of target sites under low WNT signal and can act as an activator or repressor in a context dependent manner [127–129]. This would correspond to the notion that WNT is less active in the posterior mandible, where cells are further away from the anterior WNT source and express *Axin2* at a lower level [130]. Consequently, a different WNT-induced expression profile via TCF7L2 can be generated. In addition, the transcription factor SIX2 has been shown to repress WNT targets by binding to TCF/LEF sites [129]. As SIX2 is specifically expressed in the salivary gland at E12.0, it may contribute to the suppression of WNT activity there at this stage [102].

### Patterning of the mandibular epithelium along the oral-aboral axis

In order to understand how different oral structures form in the right place and at the right time, we must first characterize how the developing mandible is patterned. By mapping the expression of region-specific markers identified from our scRNA-seq analysis at different developmental timepoints, we demonstrated that the mandibular epithelium is progressively subdivided into distinct zones along the oral-aboral axis between E9.5 and E12.0. Unexpectedly, at the onset of mandibular development almost all ectodermal cells express *Pitx2* and *Irx1*, which are dental markers at E12.0 and predicted to be the main transcription factors driving the expression of other tooth-specific genes. Notably, markers related to the further maturation and growth of the tooth germ, such as *Dsc3* and *Ntrk2*, are not expressed until after E10.5 (Fig. S10 and data not shown). This suggests that the *Pitx2*+/*Irx1*+ mandibular epithelium at E9.5 and E10.0 is transcriptionally competent of forming dental cells but has yet to initiate the full dental program. The expression of *Pitx2* and *Irx1* then become progressively confined to the forming dental lamina, as increasing numbers of anterior aboral and posterior oral epithelial cells are specified and begin to express their respective markers. Interestingly, the emergence of the posterior mandibular and tongue epithelium from a broader *Pitx2*+/*Irx1*+ epithelial band between E10.5 and E11.0 is reminiscent of the formation of discrete tooth and taste bud domains from a common *Pitx2*+/*Sox2*+ progenitor field in both Chondrichthyes (sharks) and Osteichthyes (cichlids) [131,132]. The developmental course leading to the partition of mouse dental and the more posterior epithelium may thus represent an evolutionarily basal process. Based on these data, we propose that the mandibular ectoderm is initially more dental-like, and the later specification of cells with new anterior or posterior regional identities gradually delineates the boundary of the tooth field and confines the dental lamina to its position along the oral-aboral axis. In the absence of correct patterning, as in the case of the *Tfap2a/b* mutants mentioned above, the aboral epithelium retains its dental-like identity and forms ectopic teeth [66]. Our model can also be reconciled with current ideas of tooth evolution, where the origin of oral teeth is thought to either arise from the external dermal skeleton (outside-in) or independently from the pharyngeal endoderm (inside-out) [133]. In this context, the mandibular epithelium, regardless of its ectodermal or endodermal origin, first adopted a genetic program competent of forming placodes, perhaps similar to the observed state at E9.5. The position and the final specification of the dental fate depend on how the neighboring epithelium is subsequently patterned, shifting the location of teeth or tooth-like structures along the oro-pharyngeal-aboral axis during evolution [134].

The process of epithelial regionalization we have observed in mouse mandibles is accompanied by the establishment of expression boundaries between markers labeling adjacent populations, where the overlap of gene expression gradually decreases between E10.5 and E12.0 and progressively fewer cells at the boundary co-express region-specific markers, such as *Cxcl14* and *Tfap2b* of the aboral epithelium, and *Irx1* and *Pitx2* of the dental epithelium. How these boundaries are formed and regulated in the mandibular epithelium is not understood. The juxtaposition of *Wnt7b* in the anterior epithelium and *Shh* in the dental epithelium has been proposed to determine the boundary position, as misexpression of *Wnt7a* in the entire ectoderm abolished tooth formation in cultured explants [135]. In other developing tissues with gene expression boundaries, such as the vertebrate hindbrain and the Drosophila wing, the boundary sharpness is enhanced through direct or indirect mutual repression of transcription factors downstream of morphogen-directed tissue patterning [136–138]. Such a mechanism may underlie the proximal-distal patterning of the oral epithelium during early mandibular development, as mutual antagonism between BMP4 and FGF8 delineates the presumptive incisor and molar regions respectively [10]. In this study, among the cluster-specific transcription factors we identified from the regulon analysis, several of them, including *Trps1*, *Tbx3*, *Irx1*, *Tcf7l2*, and *Six1*, have context-dependent repressor functions [107,139–141,128]. It will be informative in the future to examine their reciprocal regulation, especially as *Irx1* and *Six1* have been shown to display mutual inhibition in the *Xenopus* cranial placode [142], and IRX1 can inhibit *Trps1* expression in a murine chondrogenic cell line [143].

### NTRK2 as a regulator of early tooth morphogenesis

Our dataset serves as a useful resource to study novel regulators of epithelial morphogenesis. Using the developing tooth as a model, we were interested in understanding the functional role of *Ntrk2* because of its unique expression pattern. While previous studies have implied a role in tooth innervation, based on low resolution expression analysis of *Ntrk2*, *Bdnf*, and *Ntf4* in rats [144–147], whether NTRK2 signaling can regulate tooth morphogenesis was never addressed. Using the selective NTRK2 antagonist, ANA-12, we were able to show for the first time that signaling through NTRK2 promotes tooth invagination. In the incisor, this is in part through the ability of NTRK2 to promote epithelial cell proliferation, which is critical for generating suprabasal cells and thickening the placode [28]. However, as *Ntrk2* is also expressed in the dental mesenchyme, our experiment could not rule out the possibility that NTRK2 indirectly controls epithelial proliferation and invagination through signals from the mesenchyme. However, as *Bdnf*, encoding the ligand for NTRK2, is expressed in epithelial cells adjacent to *Ntrk2*+ cells, direct BDNF/NTRK2 signaling is likely to take place within the epithelium to drive proliferation. Interestingly, ectopic expression of *Bdnf/Ntrk2* in the taste epithelium also resulted in larger taste buds with more taste cells [148], indicative of a common effect in promoting the growth of oral epithelial organs. Finally, NTRK2 may modulate tooth morphogenesis via other mechanisms, as ANA-12 did not alter proliferation in molars. For instance, TrkB-T1, the truncated form of NTRK2 lacking the kinase domain, can signal through small Rho GTPases to regulate cell shape and migration [149,150]. Given the presence of TrkB-T1 in the dental epithelium [147], it is conceivable that NTRK2/TrkB-T1 additionally promotes cell movement to propel the invagination process.

Taken together, our results have unveiled all the epithelial cell types in the developing mandible and described the spatiotemporal distribution of key markers. This work provides a valuable resource for investigating mandibular patterning and morphogenesis and offers a transcriptional roadmap to help future derivation of different oral epithelial progenitors for tissue bioengineering and regenerative medicine.

## Acknowledgements

We thank Harrison Wang for assistance with the mouse colony; Jessica Ding from Dr. Xia Yang’s laboratory at UCLA for help with scRNA-seq analysis; Dr. Wei Du and other members of the Hu laboratory for helpful discussions. We also would like to thank Dr. Amnon Sharir, Dr. David Castillo-Azofeifa, and Dr. Ophir Klein for their comments on the manuscript. We acknowledge the UCLA Broad Stem Cell Research Center Microscopy Core for providing confocal microscopy and the UCLA Technology Center for Genomics & Bioinformatics Core for performing the sequencing. Finally, we apologize to our colleagues whose work is not cited owing to space limitations.

## Author contributions

Q.Y. and J.K.H. conceived the project, performed the experiments, analyzed the data, and wrote the manuscript. Q.Y. and A.B. carried out imaging analysis. All authors contributed to the planning of the experiments and thoroughly reviewed the manuscript.

## Funding

This work was funded by R90DE031531 to Q.Y., and R00DE025874 and R03DE030205 to J.K.H from the National Institutes of Health/National Institute of Dental and Craniofacial Research (NIH/NIDCR).

## Competing interests

The authors declare no competing or financial interests.

## Notes

### Competing Interest Statement

The authors have declared no competing interest.

